# Competitive Inhibition of Cortisol by Prostaglandins at the Ligand Binding Domain of Glucocorticoid Receptors

**DOI:** 10.1101/851501

**Authors:** Charles D. Schaper

**Author notes:** Corresponding author: Charles D. Schaper, Ph.D.

## Abstract

Cortisol is a steroid hormone that binds to the glucocorticoid receptor with a diverse range of functions including metabolic, neurologic and immune responses. Prostglandins are lipid autacoids that participate in a wide range of functions including the inflammatory response and the vasoactivity of blood vessels and smooth muscle. Previously, a direct interaction between the two classes of biomolecules has not been studied, and here it is shown for the first time that there is competitive inhibition of cortisol and prostaglandins at the critical ligand binding domain of glucocorticoid receptors, which are found in nearly every cell and tissue in the human body. To assess molecular similarity that would enable competitive inhibition, it is first noted that cortisol (CORT, C_21_H_30_O_5_) and prostaglandin E2 (PGE2, C_20_H_32_O_5_) are of nearly the same molecular weight, and have nearly the same number of carbon, hydrogen and oxygen elements. The carbon-carbon double bonds on each of the two branches of PGE2, combined with the five-carbon cyclic centering structure, enable a conformational isomer of PGE2 to position the five oxygen-based functional groups at similar spatial positions as the corresponding five oxygen-based functional groups of CORT. Within the gene-regulating ligand binding domain (LBD) of the glucocorticoid receptor (GR), the average inter-molecular spacing of hydrogen bonding to the corresponding identical five functional groups is 1.778 Å for CORT versus 1.746 Å for PGE2. The conformational isomer of PGE2 within the LBD is at a lower energy state than CORT by 54.9%, indicating preference in stability. Both PGE2 and CORT exhibit a hydrophobic core within the LBD of GR, which is compatible with hydrophobic regions of the LBD. With the incorporation of two calcium ions into both PGE2 and CORT, the energy is further reduced by 58.9% and 113% respectively. PGE2 with calcium ion stabilization is lower by 15.5% in comparison to the equivalent arrangement of CORT. Because of the steroid ring structure of CORT, the spacing of the calcium ions is larger at 16.2Å, whereas PGE2, with just a single ring, collapses to 12.3Å. Analysis of experimental data associated with the adrenal response to an injection of lipopolysaccharide (LPS) is presented to indicate competitive inhibition of PGE2 is responsible for an observed lack of phosphorylation of the GR despite the presence of CORT. Competitive inhibition preventing normative cellular processing of cortisol, a stress hormone, by prostaglandins, which are associated with the inflammatory response, may be the causative source of certain signs and symptoms of disease, and thus is of significant medical importance.

## 1 Introduction

The hormone cortisol (CORT) is released mainly from the adrenal cortex in response to adenocorticotropic hormone (ACTH), which is generated at the pituatary gland as directed by the hypothalamus to form the HPA (hypothalamic-pituatary-adrenal) axis. The target of cortisol is the ligand binding domain (LBD) of glucocorticoid receptors (GR), which are widely distributed to nearly every cell, tissue, and organ in the human body, responsible for a range of metabolic and other functions citekino2017glucocorticoid. Further, cortisol is an integral part of the limbic system which is involved in the formation of thoughts that influence motivation, fear, and mood. Moreover, cortisol is a permissive hormone inducing cells to utilize other hormones to its fullest extent, and thus, in addition to a direct effect, cortisol indirectly influences many other body functions as it enables other hormones [1].

Cortisol is a significant component of the inflammatory response as the HPA axis is activated through the immune system and the production of ACTH [2]. The influence of the prostaglandins, particularly the autacoid PGE2 which is associated with inflammation, has been analyzed in its influence on cortisol concentrations when activated, [3]. However, the particular mechanism for PGE2 in its correlation with cortisol has not been identified despite the increase in ACTH concentrations when presented with lipopolysaccharide (LPS) intraveneous injections, which also produces PGE2. Spiga and co-workers [4] utilized LPS injections to generate ACTH and CORT responses, as well as the cytokines IL-1*β*, IL-6, and TNF-*α*, but noted that there is a lack of phosphorylation of the glucocorticoid receptor at the adrenal cortex.

In this article, it is demonstrated that CORT and PGE2 exhibit competitive inhibition at the LBD of GR, in which the binding affinity for PGE2 is approximately equivalent to that of CORT. Furthermore, this result will explain and organize causality in the noted correlations of the stress hormone, cortisol, and the immune response to infection and disease involving the prostaglandins. The result is significant as there has been wide reports in the literature of linkage between cortisol and diseases, which are capable of producing PGE2: such as, in viral infection [5], neuroimmunity[6], renal cancer [7], depression [8], and somatization disorder[9]. Morever, as PGE2 is a pyrogen responsible for fever, and glucocorticoids are a noted anti-pyrogen agent, the work may enable a mechanism to describe the production of fever, which currently is lacking [10]. Thus, competitive inhibition of the normative processing of cortisol due to PGE2 binding of the LBD of GR may enable a mechanism of action to characterize the inflammatory response.

## 2 Methods

- Molecular Modeling: For the modeling of the molecules CORT and PGE2, as well as the association with calcium ions, the software program Avogadro was used. The cortisol molecule was first prepared. Then, the amino acid residues of the ligand binding domain, including the *α* carbon, were positioned in proximity to the functional groups as follows: Gln111: O3; Thr208:O1,O2; Asn33:O4; Gln39:O5; Arg80:O5. A water molecule was included about the periphery of O5. These residues were positioned in accordance with the approximate layout presented in [11], which used crystal studies for identification of the amino acid residues of the ligand binding domain of the glucocorticoid receptor. The force field MMFF94 [12] was utilized and an optimization program available in the software that minimizes the energy associated with the complex was used for determining the bond length and the positioning of the chemical elements, including the functional groups of the ligand binding domain. Two potassium ions were added in the vicinity of high negative electrostatic potential and the optimization program utilized to minimize the energy in positioning the potassium ions. This process was repeated for calcium ions. The dimensions of the spacings were determined at the central location of the carbon, oxygen, or hydrogen element, depending upon whether it was the size of the ligand binding domain or the length of the hydrogen bonding. The electrostatic potential was computed using the software. After determining the positional configuration of cortisol and the amino acid side chains comprising the ligand binding domain, the cortisol molecule was removed while retaining the positioning of the LBD amino acid residues. The water molecule was removed. The PGE2 molecule then was inserted into the LBD in a conformer state as follows: Gln111: O3; Thr208: O1, O2; Asn33: O4; Gln39: O5; Arg80: O5. A water molecule was added in the vicinity of O3. The optimization software program was then run to determine the optimal bond lengths and positioning of the elements. Potassium ions and calcium ions were added in the vicinity of high negative electrostatic potential and the software program was run to determine the optimal position. Hydrogen bonding length and the positioning of the alpha carbon of the amino acid residues comprising the ligand binding domain were measured. The electrostatic potential was then calculated. Placement of a third calcium ion was attempted in an area of moderate negative electrostatic potential, and the optimization program was run, which resulted in the calcium ion moving away from the molecular complex, indicating that the complex could only hold two calcium ions, and not three.
- Molecule Selection: The competitive inhibition of CORT by PGE2 was evaluated because of their potential for matching in chemical elements, molecular weight, hydrogen bonding, core cyclic structures and hydrophobic characteristics, electrostatic interaction, intramolecular interactions, and topography, while retaining the basic structural characteristics, such as the *cis* and *trans* positioning structure of double bonds to achieve the approximate comparative structure of Table 1. Moreover, a prior study conducted by this lab indicated the potential for interaction of cortisol and prostaglandins in producing fevers and symptoms of other diseases [10], and a molecular basis for this association was sought. The supplemental information provides the code and the videos of the three-dimensional arrangement of CORT and PGE2 conformer at the residues of the amino acids comprising the functional LBD of GR.
- Experimental Analysis: To demonstrate experimentally the competitive inhibition of cortisol by prostaglandins at the LBD of GR, the experimental data and analysis of Spiga and co-workers [4, 13] was utilized. In this case, experimental results were produced for the situation where an injection of lipopolysaccharide (LPS) was administered intravenously to trigger the response of the adrenal gland. Measurements were taken over time that included adrenocorticotropic hormone (ACTH), cortisol, TNF-*α*, IL-1*β*, IL-6, and the phosphorylation of GR (pGR) to indicate its activity. In this experimental set, pGR was near zero, indicating a lack of GR activity, whereas a prior experiment with ACTH injected directly, pGR activity was present. For both experiments, CORT concentration was present and similar. In Spiga et al.’s analysis [4], a mathematical model was developed to characterize the lack of activity of GR. Activity in GR would have been present because of the high levels of CORT due to ACTH presentation at the adrenal gland, which should have expressed GR activity, but which did not. Thus, [4] revised the model to indicate that TNF-*α* inhibited the interaction of CORT with GR, and thus a mathematical function was applied that enabled a better fit to the data. In the developments of this article, it is indicated that while TNF-*α* plays a role, it is rather the competitive inhibition of PGE2 on GR that explains the lack of phosphorylation of GR in this experimental set of data. As described in the supplementary information for this article, the model equation 23 of [4] supplemental was updated to now include Michaelis-Menten dynamics for competitive inhibition of PGE2. This expression was then used to compute the PGE2 concentration to match both the experimental data and the function *φ*_*pGR*_(*TNFα*) of the revised model described in Section 3.1,” Model Equations for the SRN with Cytokine Interactions” of [4]. The calculations are presented in the supplementary information of this paper, Section 1.

## 3 Results

To further analyze the potential for CORT and PGE2 to compete for association with the LBD of GR, the positional distribution of the respective functional groups are analyzed. In an important step in the comparison, conformational isomers, or conformers, of PGE2 are evaluated through wireframe sketches to assess a feasible configuration that is representative of the functionality of CORT. Detailed molecular models of the most promising conformer of PGE2 is then compared to CORT to determine the bond length, positioning, and spacing of the intramolecular functional groups. The integration of the molecules and the functional groups at the LBD of GR are computed in terms of intermolecular spacing, hydrogen bonding, electrostatic potential, molecular energy, and intramolecular characteristics. The energy levels and the configuration in association with calcium ions are evaluated in terms of its electrostatic potential. A pathway is developed for competitive inhibition, and it is evaluated by comparison with published experimental data.

**Table 1:** Initial comparison of the chemical affinity of Cortisol and a conformer of Prostaglandin E2 at the ligand binding domain of glucocorticoid receptors, before molecular modeling.

| Item | Cortisol | Prostaglandin E2 Conformer |
| --- | --- | --- |
| Molecular Weight | 362.46 g/mol | 352.46 g/mol |
| C elements | 21 | 20 |
| H elements | 30 | 32 |
| O elements | 5 | 5 |
| core | hydrophobic | hydrophobic |
| core C | 6 | 6 |
| periphery binding | hydrogen bonding | hydrogen bonding |
| periphery groups | three O/OH | three O/OH |
| O at (I/II/III) groups | 3/1/1 | 3/1/1 |
| O5 functional group | O + H<sub>2</sub>O | OH |
| O3 functional group | OH | O + H<sub>2</sub>O |
| Approximate Size C-C | 8 x 5 | 8 x 5 |

### 3.1 Comparing CORT and PGE2

In Figure 1, the chemical structure of CORT and PGE2 is presented in the standard arrangements. To make the case, it is first noted that CORT (C_21_H_30_O_5_) and PGE2 (C_20_H_32_O_5_) are of similar molecular weight, 362.46 g/mol versus 352.46 g/mol respectively, and have nearly the same number of chemical elements: 21 versus 20 carbon; 32 versus 30 hydrogen; and importantly the same number of oxygen elements, five, which can be arranged as five functional groups. When expressed in standard format the overall structure of PGE2 is dissimilar to that of the structure of CORT; however, through single bond rotations at C4 and C12 of PGE2, a structure is formed resembling closer to that of CORT. This structural expression maintains the same double bond arrangement of *cis* configuration at C5-C6 and *trans* configuration at C13-C14, and thus is the same molecule. The arrangement critically positions three oxygen groups together such that it is comparable in position to that of CORT at C1, C2 and C3. With further single-bond rotation at C15, a match of the functional groups is achieved, and with rotation at C16, C17, the dimensional size of CORT can be approximated by PGE2. Further, lateral stability is established through the interaction of C20 with the double bond of C5-C6, and a hydrophobic core is achieved, thus matching well the cyclic structure of CORT. Further, it is noted that the five functional groups of the conformer of PGE2 are all positioned on the periphery, as in CORT, and the core is hydrophobic, as is CORT. There are groups of three functional groups located on one side of the molecule for each, that is O1, O2, and O3 are approximately equivalent, and there is one oxygen group in the middle for each, and one oxygen group on the opposite side. Along one edge for both molecules, no functional groups are present.

**Figure 1:**
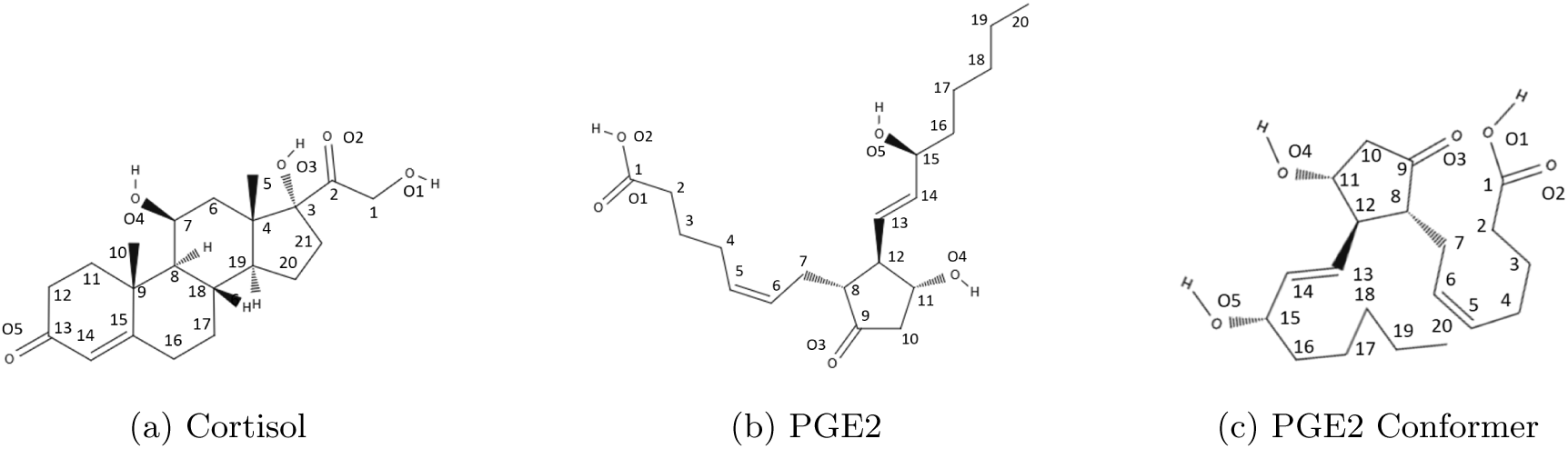
*A comparison of the molecular arrangements of (a) cortisol*, C_21_H_30_O_5_ *and (b) PGE2*, C_20_H_32_O_5_ *in nominal configuration. In (c) a conformer of PGE2 is shown, that more closely resembles the cyclic structure of cortisol. Note that the cis and trans configurations of the carbon-carbon double bonds are conserved in the conformer of PGE2 relative to PGE2, and thus are the same molecule but just organized in a different positional arrangement.*

### 3.2 Structural Analysis

To evaluate the positional relations and dimensional similarity of the functional groups comprising CORT and PGE2, three dimensional models are developed using the molecular simulator, employing a standard force field of MMFF94. In Table 2, the intramolecular distances are compared between the five oxygen-based functional groups of CORT and those corresponding to the conformer of PGE2: In general good agreement is indicated between CORT and PGE2, within the three functional group regions. Especially important, is that for the five membered carbon ring of PGE2, which corresponds to the spacing of O4-O3, it is noted that it is in good agreement with the distance of O4-O3 of CORT. This is important not only because it connects the I and II functional groups, but also because the distance for O4-O3 of PGE2 is fixed due to the cyclic structure. Whereas, the O5-O4 and the O3-O2 groups of PGE2 can be adjusted to optimal distances because of the freedom of motion due to the two branches. It is noted also that the *trans* and *cis* configurations of PGE2 aid in the positioning of the functional groups of the structure, and will provide stability and increase the probability of association with the corresponding binding sites. The distance of O3-O2 is different between CORT and PGE2. However, one intermediary water molecule will be introduced to CORT and to PGE2, but at different locations, that will establish an equivalent configuration for hydrogen bonding. Moreover, the PGE2 oxygen based groups corresponding to the carboxylic acid group on a largely unconstrained branch of PGE2 will enable optimal positioning. The projected dimension is approximately 11.4Å *×* 5.0Å for CORT and 10.6Å *×* 4.7Å for PGE2. Thus, both CORT and PGE2 conformer have the same aspect ratio of 2.3 before insertion into the LBD.

**Table 2:**
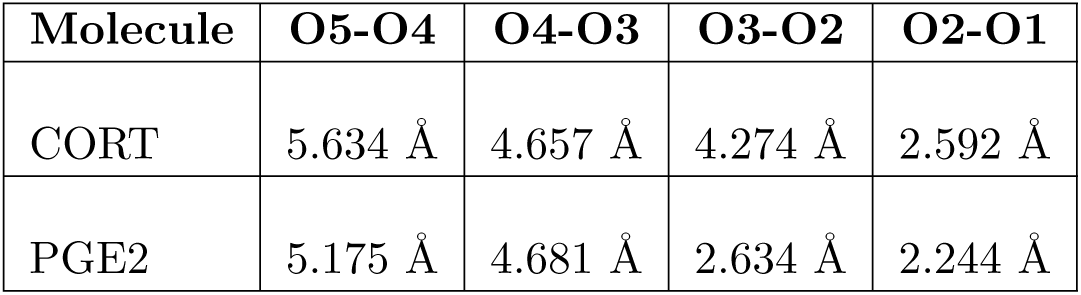
The distance between the five functional groups of cortisol and a conformer of prostaglandin E2 is compared. The five membered ring of PGE2 corresponds to the critical O4-O3 group, and it is noted that it is in agreement with that of CORT. This is critical because it connects the I and II functional groups, and the distance for O4-O3 PGE2 is fixed due to the ring structure; whereas the O5-O4 and O3-O2 groups of PGE2 can be adjusted to optimal distances by motion of the branches. The distance of O3-O2 is different between CORT and PGE2. However, the PGE2 group corresponds to the carboxylic acid group and because of its electrostatic potential may provide equivalent binding capability as the hydroxyl group of CORT despite the longer difference to the corresponding attachment site at GR.

| Molecule | O5-O4 | O4-O3 | O3-O2 | O2-O1 |
| --- | --- | --- | --- | --- |
| CORT | 5.634 Å | 4.657 Å | 4.274 Å | 2.592 Å |
| PGE2 | 5.175 Å | 4.681 Å | 2.634 Å | 2.244 Å |

In Figure 2 the volume of space occupied by the molecules are compared by depicting the van der Waals hard shell radius for CORT and the conformer PGE2 under study. It is seen that the volume and positional arrangement of the functional groups is similar. The periphery space occupied by the three functional regions have similar presentation, and are accessible to intermolecular forces which will be important for the PGE2 conformer to associate with the functional groups of the LBD. The relative dimensional profile is similar and the hydrophobic nature within the core region is similar.

**Figure 2:**
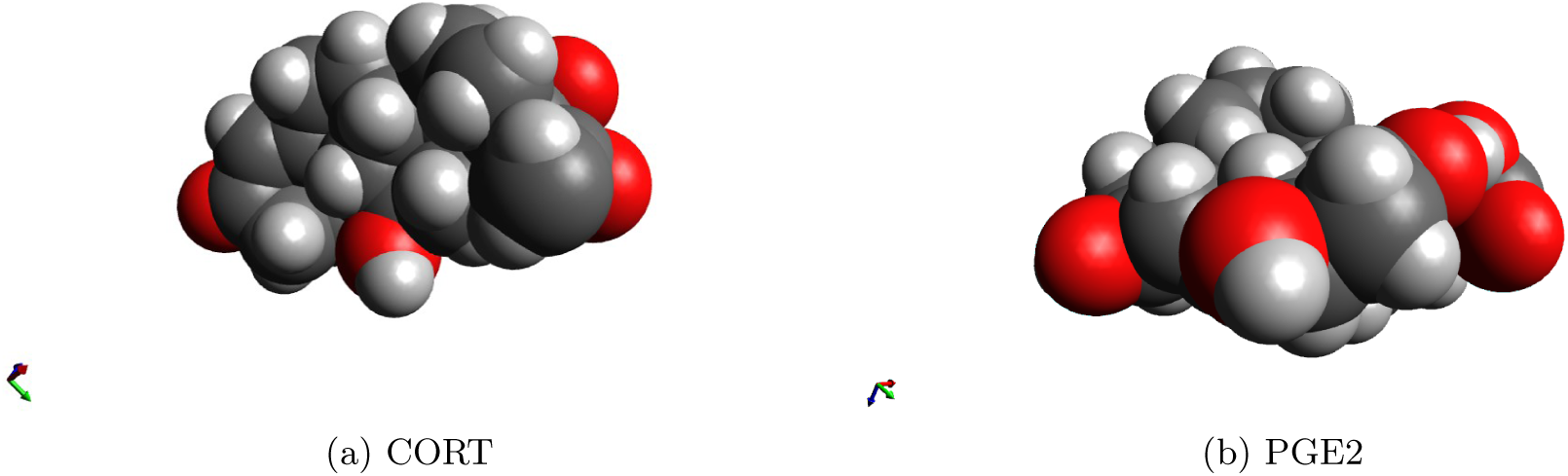
For cortisol and prostaglandin E2, the van der Waals hard shell configurations indicates the peripheral positioning of the functional groups, which are thus accessible for binding to the corresponding areas of the LBD of GR.

### 3.3 Ligand Binding Domain of Glucocorticoid Receptors

Recently, high resolution crystal structures of the ligand binding domain of glucocorticoid receptors have identified the amino acid groups responsible for the positioning of cortisol [11]. In this section, the spatial relationship of PGE2 conformer is developed based upon the amino acids identified in these crystal structures, and compared against cortisol. We first analyze simple wire sketches, which are illustrative, and then apply the molecular modeling software programs for spatial analysis.

In Figure 3, the spatial positional relationship for a wireframe approximation is presented of CORT and PGE2 within the LBD of GR, to indicate the potential for similar configurations of the functional groups with respect to the functional groups comprising the LBD. It is noted that the functional groups PGE2-GR is appropriate to be placed in three regions. It is further noted that the conformer of PGE2 may even be preferred in comparison to CORT-GR as III_CORT_ requires the presence of water (not shown), whereas with III_PGE2_ the hydroxyl group is available to interact directly with Gln39 and Arg80. In I_PGE2_ the single hydroxyl group is available for interaction with Gln111 and Thr208. Further examination of Figure 3 also indicates the importance of the five-membered carbon ring, as it positions the two branches of PGE2 to associate with the hydrogen bonding centers of LBD of GR. This establishes the hydrophobic core including the potential multi-ring structure, and the functional groups surrounding the core. Moreover, the ring structure of PGE2 sets-up the functional groups on the exterior of the molecular and an internal hydrophobic core, which includes a pair of carbon-carbon double bonds at C5-C6 and C13-C14 interacts with groups of GR including Leu77, Met73, Trp69, Met70, Leu32, Phe92, Cys205, Phe204 (not shown in figure but positioned above and below the internal core structure of PGE2).

**Figure 3:**
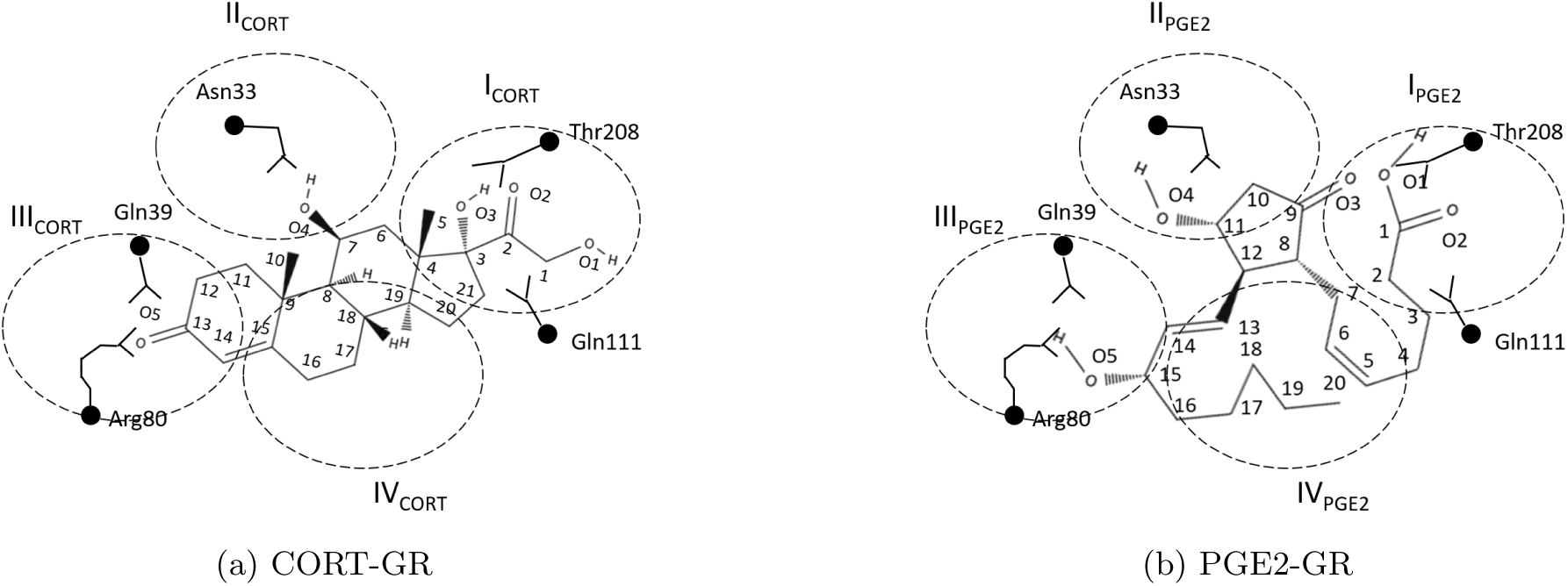
For the ancestral glucocorticoid protein receptor (AncGR2), which is a stable GR useful for crystal studies, at the ligand binding domain, a fit of a conformer of PGE2 very similar to that of CORT is noted. There is possible equivalent hydrogen bonding at the three functional groups, and at the interior there is substantial hydrophobic interaction and similarity. There may be an improved interaction with PGE2 relative to cortisol with Gln39 and Arg80 since CORT requires an external water molecule (not shown) since only a carbonyl group is available, whereas PGE2 already has a hydroxyl group. The three membered oxygen group of PGE2 contains enough free motion as it is unconstrained by rings to achieve optimal alignment. The double bond alignment of PGE2 at the hydrophobic core is noted. It is also noted that the ring structure of PGE2 formed via the C8-C12 bond is critical to achieve the proper configuration of the two branches and subsequent fit within the hydrophobic core, which incorporates two carbon-carbon double bonds.

To assess the three dimensional nature of the intermolecular arrangements, the molecular modeling program is used to optimize bond lengths, elemental positioning, and spacing of the functional groups. The three dimensional positioning of CORT and PGE2 as a conformer is exhibited in Figure 4. As indicated earlier, the water molecule for CORT is positioned between O5 and Arg80, while for PGE2, the water molecule is positioned between O3 and Thr208. Both show similar interaction with the binding regions of the GR protein, in terms of intermolecular spacing. The program converges to a local minimum as the positioning initially of the molecules influences the ultimate convergence, including with respect to the position of the hydrogen groups. The van der Waals space filling model representations are also presented for CORT and PGE2 to show that both molecules occupy similar volume within the LBD. The arrangement of the functional groups along the periphery show similar distance spacing. One of the branches of PGE2 does show out of plane positioning compared to the tighter profile of CORT.

**Figure 4:**
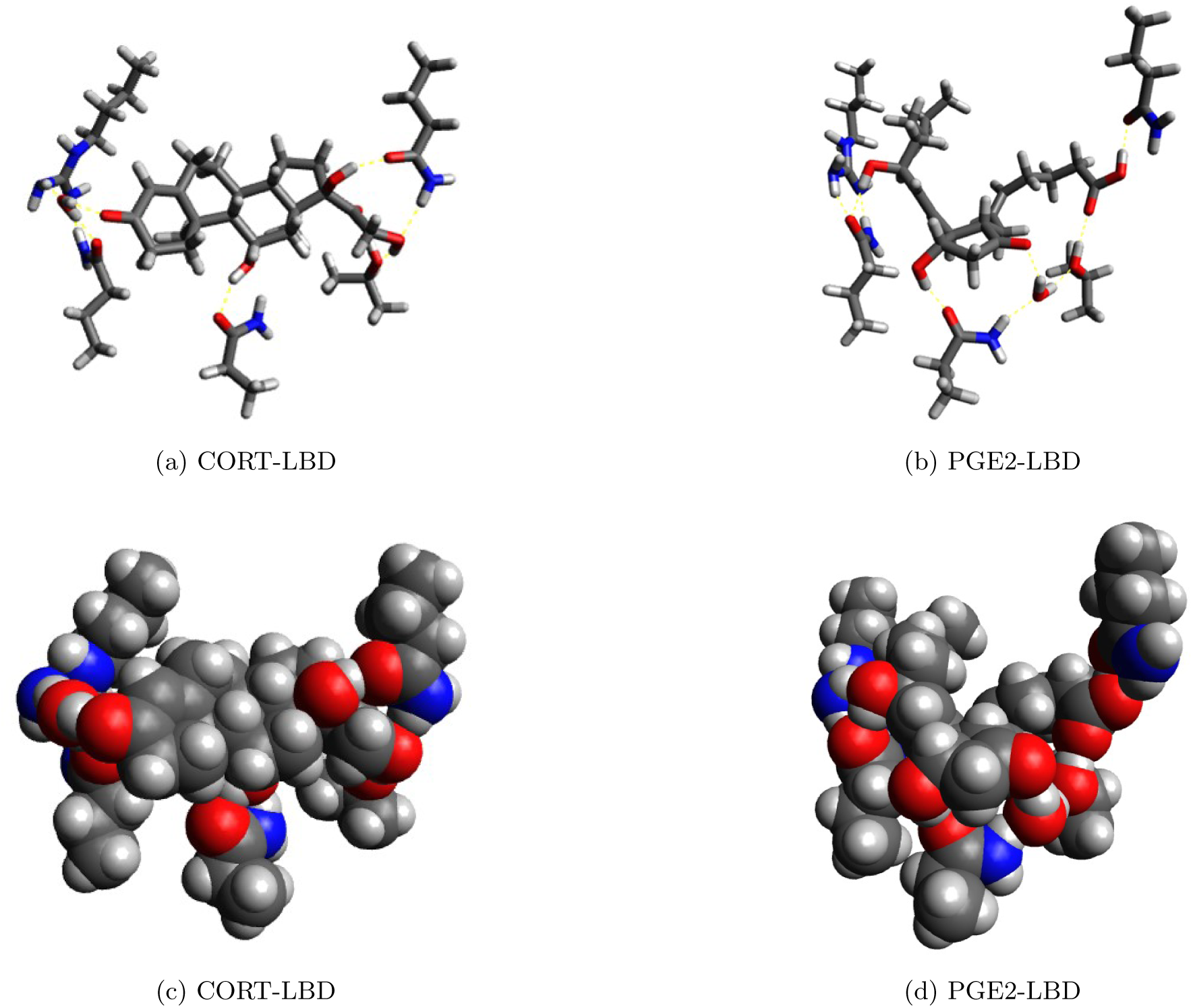
For representations of the positional arrangement and space-filling molecular models, the arrangements are depicted within the LBD of the GR for (a) and (c) cortisol and (b) and (d) PGE2 conformer. The free branch of PGE2 is pushing one of the amino acid groups Arg80 outward relative to that of CORT. Significant hydrogen bonding is noted for both CORT and PGE2.

Related to the molecular dynamics of introducing the PGE2 molecule into the slot at the LBD of GR which nominally is designated for CORT, the dimensions were calculated of the PGE2 molecule with rotation at C12 but prior to the final conformational arrangement involving the positioning of the carboxylic acid group. The linear structure of PGE2 positions the three leading functional groups with a spacing of 4.627 Å (O5 to O4) and 4.640 Å (O4 to O3) prior to the final configuration. This indicates a positional arrangement that is nearly the same in dimension, thereby providing the possibility of the orientation of PGE2 being reversed when presenting to the LBD of GR; that is, for example, O3 may associate with Gln39 and ARG80. Also, it is noted that the five-membered ring and the double bond carbon achieves a planar structure consistent with the nominal corticosteroid structure of CORT, and enabling its entry. Further it is noted that the spacing to the carboxylic group from the lead functional groups to the double bond on the alpha group is 6.569 Å, which is greater than the width of the LBD structure in the final configuration. Hence, there is sufficient size for PGE2 to enter without the possibility for out-of-plane steric hinderance from the upper and lower portions of the slot opening of LBD of GR. Thus for the molecular dynamics of positioning of PGE2, it seems likely that the five carbon ring is first positioned at I_PGE2_ and II_PGE2_. The hydrophobic core then drives the hydroxyl group on the second (*β*) branch to III_PGE2_ and then the carboxylic acid group of the first (*α*) branch to bind to the associated LBD corresponding to the functional groups of I_PGE2_. The C20 carbon is then positioned out-of-plane with respect to the double bond groups of C5 and C6 to produce a low energy state.

In terms of intermolecular spacing of the five oxygen-based functional groups from the corresponding amino acid side chains, Table 3 provides a comparison of the hydrogen bonding of CORT and PGE2 within the LBD of GR. The range of spacing for CORT is from 1.720 Å to 1.832 Å, and for PGE2 the range is from 1.598 Å to 1.868 Å; the average spacing of the hydrogen bonding for CORT is 1.778 Å and for PGE2, the average spacing is 1.746 Å. The water molecule for PGE2 was placed in the vicinity of O3 and Thr208. The spacing from the inserted water molecule of PGE2 at O3 to Thr208 is 2.761 Å; for CORT, the spacing of the oxygen to the the oxygen element comprising the water molecule to Arg80 at O5 is 2.732 Å. For the configuration of PGE2, the average bond distance for the nearest neighbor is 1.760 Å, which is tighter than CORT by 1.842%. In fact, each oxygen group has hydrogen bonding attraction to a side chain group or the inserted water group. For all, the carboxylic acid area has the shortest intermolecular distance at 1.598 Å.

**Table 3:**
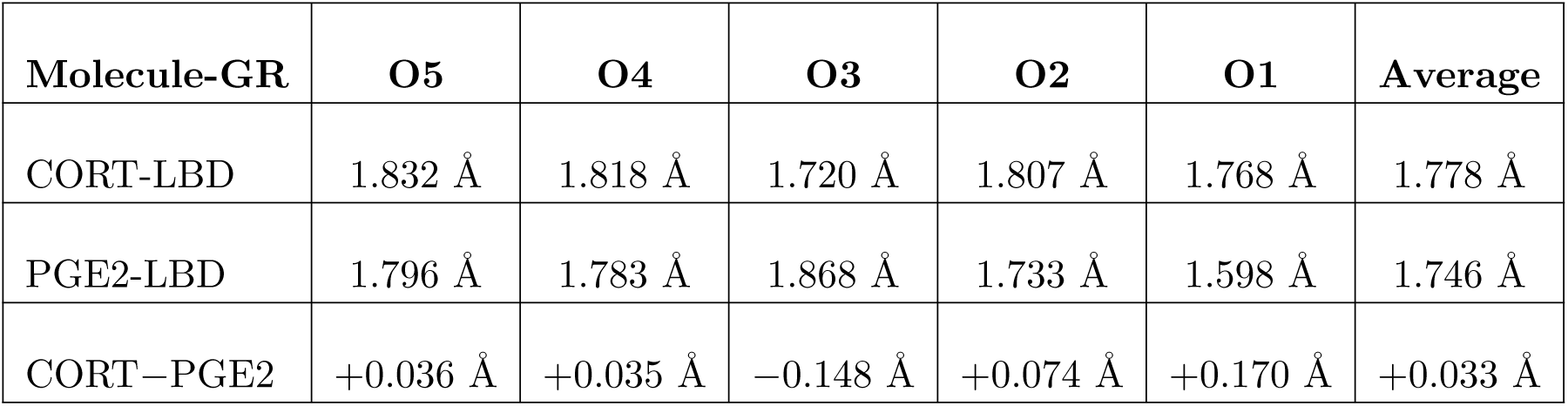
*The intermolecular spacing of the oxygen-based functional groups to amino acid side group elements comprising the LBD. For CORT, the O5 group space is to a water molecule which is then associated with Arg80, and for PGE2 to O3 group space is to a water molecule which is then associated with Thr208. The average spacing for CORT is calculated at 1.778* Å *and for PGE2, the average spacing is 1.746* Å. *The average difference in spacing is thus 0.033* AA.

### 3.4 Incorporating Calcium Ions

It has been indicated that there is an association of two calcium ions when cortisol enters the cytosol [14], while the actual reasoning or usage of calcium with cortisol is not indicated. In this section, the potential role of calcium is explored for both CORT and PGE2 at the LBD of GR to show that both molecular structures have two sites each where calcium ions can be incorporated; we will later evaluate the energy of the overall structure in section 3.7. For the introduction of calcium ions, the molecular organization and a comparison is indicated in Figure 5(c) for CORT and in Figure 5(d) for PGE2. Similarity is noted in terms of the incorporation of the calcium ions at two ends of the molecular arrangements, which disrupts the hydrogen bonding, and positions the calcium ion within the central area for electron sharing with three or four of the oxygen based functional groups. It was not possible to incorporate a third calcium ion at other regions of the overall structure.

**Figure 5:**
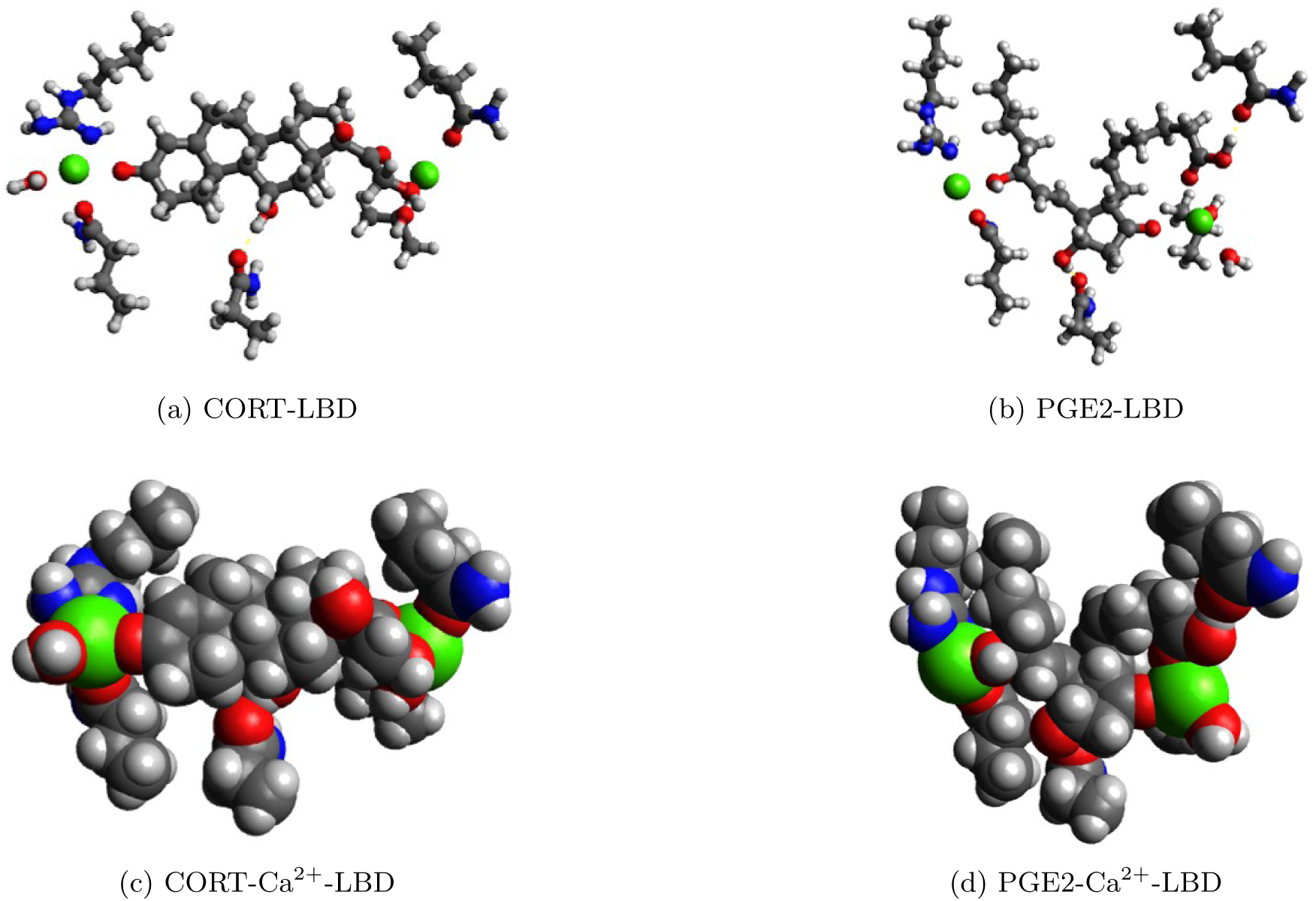
With two calcium ions optimal positioned within the CORT-GR complex (a) and (c), it is noted that the oxygen based functional groups of CORT come into association with Calcium ions at both ends of the molecular arrangements, which is similar as that of PGE2 (b) and (d). It is noted that the spacing of the calcium ions for CORT is greater than that of PGE2, since CORT has four cyclic structures, thus preserving the structural arrangement as that affiliated solely with hydrogen bonding, whereas the collapse of PGE2 is due to the single cyclic structure and one double bond.

### 3.5 Structural Comparison

To evaluate the potential motion of the *α* carbons of the amino acid residuals comprising the LBD, the distances were calculated and compared for the introduction of CORT and of PGE2. In Table 4, the distances between the *α* carbons of each of the functional groups of the LBD is calculated, along with the differences. In general, the intermolecular spacing of the groups comprising the GR are similar for CORT and PGE2, as the average distance for the LBD is between 13.047 and 13.840 Å, within a min-max of 5.919% to the mean, for each of the configurations indicating that the LBD shifts the relative positioning of the groups to accommodate the different molecules.

**Table 4:** *Difference in position of the α carbon of the amino acid side chains comprising the functional groups of the LBD of GR. It indicates that while the functional groups comprising the attachment points to either CORT and PGE2 are positioned almost identical, the α carbons comprising the protein structure are relatively stable. The largest differences are seen when calcium ions are incorporated into the structures. (C-Ca denotes CORT with two Ca*^2+^ *configuration; P-Ca PGE2-Ca*^2+^; *ΔC denotes the difference in spacing of CORT and CORT-Ca*^2+^; *ΔP denotes the difference in spacing of PGE2 and PGE2-Ca*^2+^; *P C denotes the difference in spacing of PGE2 and CORT; ΔPC denotes the difference in spacing between CORT and PGE2 with Ca*^2+^ *ions present.)*

| Cα spacing | | CORT | PGE2 | C-Ca | P-Ca | $\Delta C$ | $\Delta P$ | P-C | $\Delta PC$ |
| --- | --- | --- | --- | --- | --- | --- | --- | --- | --- |
| Thr208 | Gln111 | 11.523 | 13.472 | 10.389 | 12.365 | -1.134 | -1.107 | 1.949 | 1.976 |
| Thr208 | Asn33 | 7.770 | 8.029 | 9.681 | 6.766 | 1.911 | -1.263 | 0.259 | -2.915 |
| Thr208 | Gln39 | 13.306 | 10.717 | 14.537 | 10.191 | 1.231 | -0.526 | -2.589 | -4.346 |
| Thr208 | Arg80 | 15.921 | 17.436 | 16.484 | 18.242 | 0.563 | 0.806 | 1.515 | 1.758 |
| Asn33 | Gln111 | 15.886 | 17.942 | 14.823 | 16.400 | -1.063 | -1.542 | 2.056 | 1.577 |
| Asn33 | Gln39 | 7.100 | 5.262 | 6.400 | 5.381 | -0.700 | 0.119 | -1.838 | -1.019 |
| Asn33 | Arg80 | 15.508 | 17.780 | 14.874 | 18.750 | -0.634 | 0.970 | 2.272 | 3.876 |
| Gln39 | Gln111 | 18.975 | 18.530 | 17.941 | 17.179 | -1.034 | -1.351 | -0.445 | -0.762 |
| Gln39 | Arg80 | 12.950 | 13.959 | 12.790 | 15.147 | -0.160 | 1.188 | 1.009 | 2.357 |
| Arg80 | Gln111 | 13.019 | 15.272 | 12.554 | 14.665 | -0.465 | -0.607 | 2.253 | 2.111 |
| Average |  | 13.196 | 13.840 | 13.047 | 13.509 | -0.148 | -0.331 | 0.644 | 0.461 |
| Standard Deviation |  | 3.695 | 4.553 | 3.454 | 4.709 | 1.048 | 1.032 | 1.757 | 2.612 |
| Range |  | 11.875 | 13.268 | 11.541 | 13.369 | 3.045 | 2.730 | 4.861 | 8.222 |

The most significant difference corresponded to the introduction of the calcium ions. Because CORT contains four ring structures while PGE2 contains only one, a significant discrepancy existed between the placement of calcium ions, which were separated by a distance of 16.279 Å for CORT while only 12.366 Å for PGE2. In addition the largest difference in the positioning of the LBD was the most for calcium introduction at 8.222 Å, whereas without calcium, the difference is 4.861 Å. The non-expression of GR to molecules other than CORT may be explained by the size differences associated with calcium, that is because CORT can tolerate calcium without a reduction in size, it may be beneficial to the specificity offered by the steroid hormone molecule.

### 3.6 Electrostatic Potential

The electrostatic potential is indicated in Figure 6. Similarity is noted, with the periphery groups possessing the electrostatic charge, with the central areas neutral, including one side of the structure. The areas of greatest electrostatic concentration are maintained in a comparison of the two molecules. For the introduction of calcium ions however, a significant difference is noted in the relative location of the two sites of strong electrostatic potential, because of the differences in the positioning of the calcium ions. It is suggested that the electrostatic potential offers an interactive site that will associate with nuclear translocation and ultimately with gene expression.

**Figure 6:**
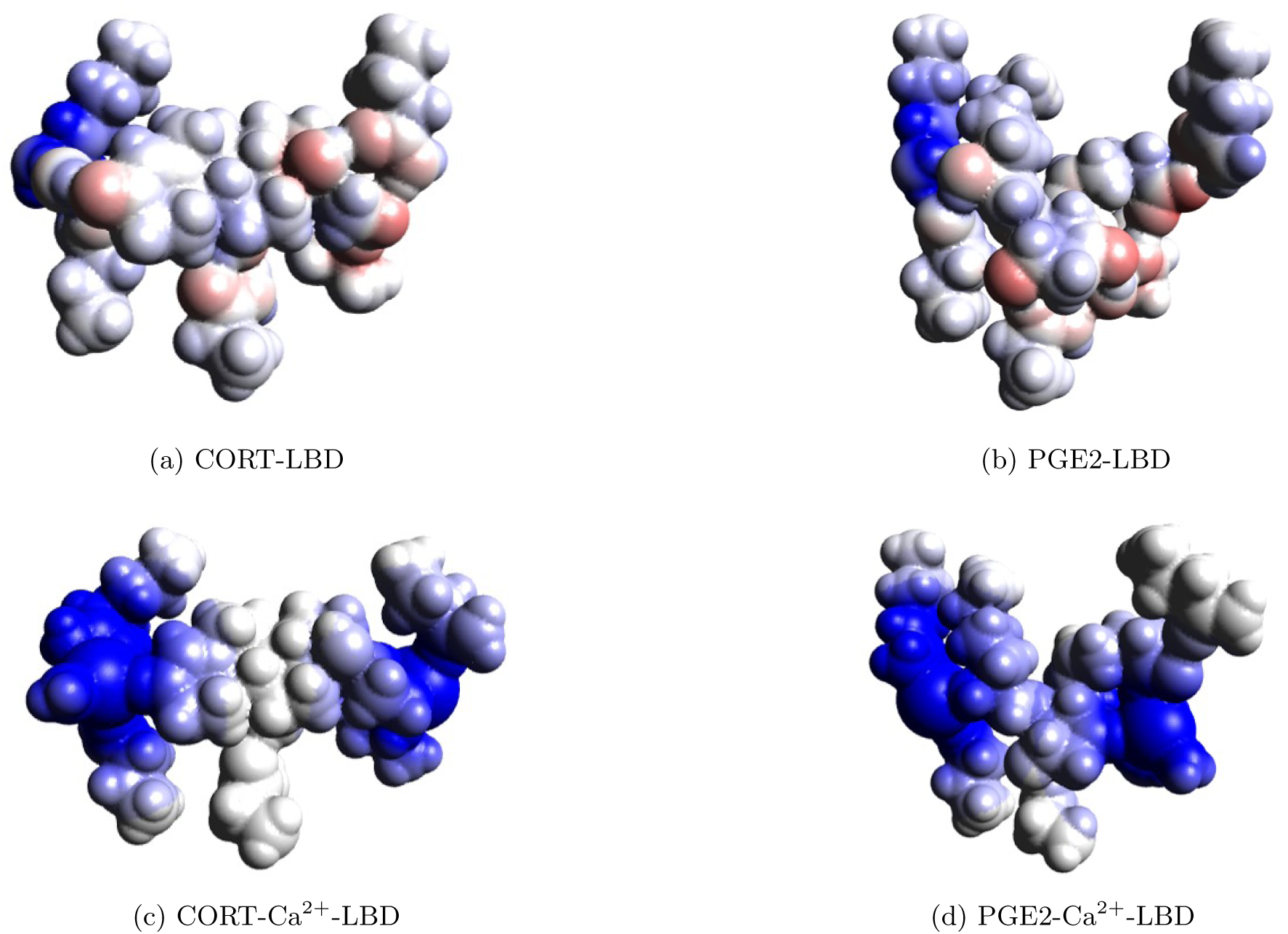
The electrostatic potential of (a) cortisol and with calcium ions added (c) and the corresponding structures (b) and (d) of PGE2 conformer. Note that with the calcium ions added, the complex becomes more attractive towards negative charge, which may permit nuclear translocation.

### 3.7 Energy Levels

The energy levels are indicated in Table 5 for the configurations. For CORT-LBD, the energy of the complex is *-*885 kJ/mol, which is actually greater than that of the LBD itself, whereas PGE2-LBD is *-*1,372 kJ/mol, which is lower than the LBD, thus more favorable. With the introduction of the calcium ions, a significant shift in the energy levels ensues, with CORT-Ca^2+^-LBD stabilizing by 113% to *-*1,888 kJ/mol., whereas PGE2-Ca^2+^-LBD is reduced by 58% to *-*2,181 kJ/mol, which is 15% lower than CORT.

**Table 5:** The energy requirements to disassociate the structures indicates that the PGE2 structures are relatively more stable than those of CORT. Moreover, the stabilizing influence of calcium ions is shown.

| Structure | Energy (kJ/mol) |
| --- | --- |
| LBD | −1,083.61 |
| CORT-LBD | −885.826 |
| PGE2-LBD | −1,372.58 |
| CORT-Ca<sup>2+</sup>-LBD | −1,888.51 |
| PGE2-Ca<sup>2+</sup>-LBD | −2,181.51 |

### 3.8 Pathway of Competitive Inhibition

In Figure 7, a basic relation describing the sequence from the introduction of active biomaterials to the resultant output of the glucocorticoid receptor, designated as *U*, is indicated. It begins with the Arachidonic acid, *A*, pathways to generate PGE2, *P*, via the cyclooxygenase enzyme, *E*. After PGE2 associates with the glucocorticoid receptor, *G*, it is in competition with CORT, *C*. The rates of the process will thus be influenced by the stability of PGE2 and CORT with GR, and based on the modeling results, would slightly favor PGE2. Additional analysis can be performed using a similar flow can be used for other prostaglandins, and for related materials to CORT, including synthetic materials, such as dexamethasone and hydrocortisone. Using this relation, mathematical models can be developed of the kinetics, which can be incorporated within overall material balances to predict the concentration variations and expressions over time.

**Figure 7:**
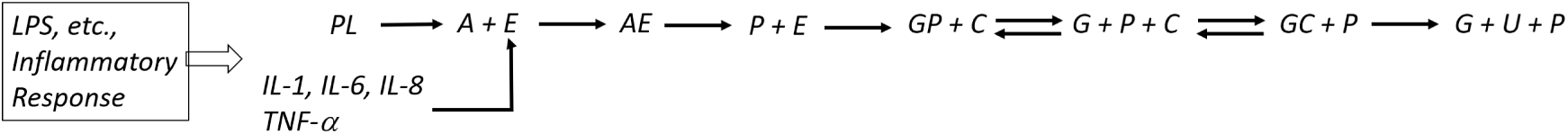
Relationship indicating the source of biomaterials, conversion to PGE2, and subsequent competitive inhibition with Cortisol at the GR. The source starts from Arachidonic Acid, which can come from the phospholipid bilayer or infectious agents, and the enzyme cyclooxygenase, which can be generated from the cytokines, for example. Either one in excess will produce PGE2, which then competes with CORT for the LBD. After association with CORT for sufficient time, the GR produces a controlled output U, such as mRNA in synthesizing new protein structures.

### 3.9 Experimental Evaluation

To evaluate the thesis of competitive inhibition, in Figure 8, a comparison of PGE2 interference at the GR is made in the experimental data of [4], in which LPS was used to trigger an ACTH response, inducing CORT from the adrenal gland. However, there was a lack of phosphorylation at the GR, which [4] indicated was due to the inhibition of TNF-*α* on GR. Here it is indicated that a more precise reason for the inhibition was due to PGE2 inhibition of the LBD at GR. According to the pathway of Figure 7, TNF-*α* increased the concentration of the enzyme COX while the LPS increased the concentration of AA, which together formed PGE2 that interacted with the LBD of GR, preventing the phosphorylation of GR despite the synthesis and presentation of CORT. The result indicates a rapid rise in PGE2, followed by a reduction and stabilization. The results indicate the concentration of PGE2 within the adrenal gland.

**Figure 8:**
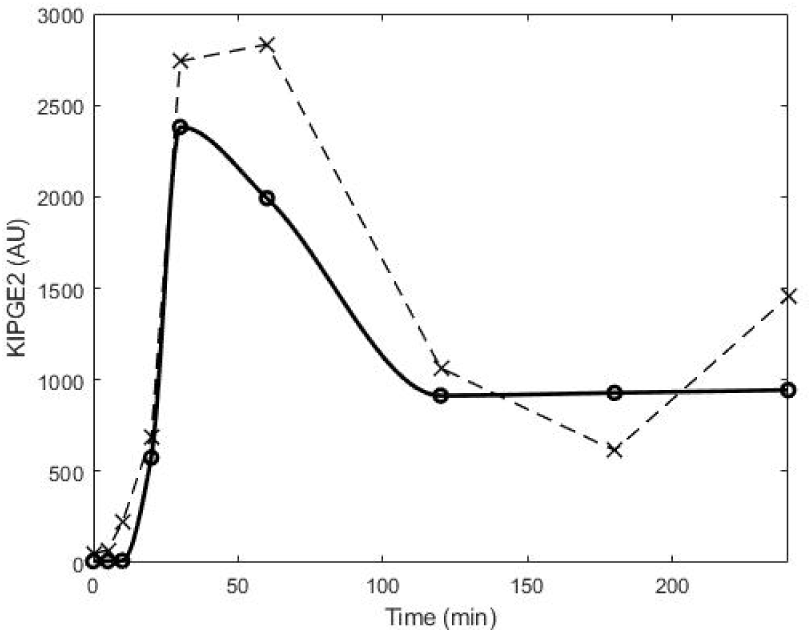
The capacity for competitive inhibition of the LBD at GR of CORT by PGE2 is evaluated for the experimental data of [4] in which a dose of LPS applied to the adrenal gland noted that there was lack of phosphorylation of GR, indicating a lack of activity. The model as described in the supplemental information is used to compute the product of a rate constant and the concentration of PGE2 necessary to explain the experimentental data, and to match the function of TNF-α that was needed in order to match the mathematical model to the experimental data.

## 4 Discussion

In this study, the competitive inhibition of cortisol by prostaglandins, particularly PGE2, has been shown for the critical ligand binding domain of glucocorticoid receptors. To explain the interference of PGE2 at the LBD of GR, a conformational isomer was indicated in which the five oxygen-based functional groups of PGE2 were aligned with the LBD sites which in normative circumstances would align with the oxygen-based functional groups CORT. The single cyclic structure and the double bonds on each of the two branches of PGE2 enable the entrance of the molecule within the LBD zone of GR and produce initial coupling with the corresponding functional groups of the LBD through hydrogen bonding. Favorable agreement for competitive inhibition in the moleculear configurations of PGE2 and CORT is apparent across multiple areas of comparison including molecular weight, number of chemical elements, relative positioning of functional groups, hydrogen bonding, hydrophobic centering, topology, dimensional size, and both intramolecular and intermolecular arrangement of structures. Moreover, the conformational isomer of PGE2 at the LBD of GR appears to be relatively stable, that is energetically lower by 15%, in comparison to CORT, and thus would constitute significant interference in the processing of cortisol.

There are structural and electrostatic potential differences between CORT and PGE2, most notably with the integration of calcium ions (which is itself a novel insight), which disable the functionality of GR when binded with PGE2. Besides the reduction in the distance between the calcium ions for the two configurations, another difference was that although one water molecule was each needed for CORT and for PGE2 to achieve the hydrogen bonding interaction, it was situated at different positions within the LBD, with the water molecule associating with Arg80 for CORT and associating with Thr208 for PGE2. There is stronger hydrogen bonding in the PGE2 configuration than that of CORT at the carboxylic acid group.

The analysis to experimental data was evident in characterizing the influence of inhibition due to PGE2 as responsible for the observed lack of phosphorylation that would have been present had CORT immediately interacted with GR. The shape of the interaction of PGE2 with the GR indicates an influx of PGE2, which would be the result of a materials comprised of the precursors to AA and COX, as well as the cytokines. A decrease in concentration of PGE2 results after the initial influx, likely due to the diffusion of the material outside of the cell as the influence of cortisol. A stabilization period follows after two hours based on the modeled response, however it may be the case that as IL-1*β* and IL-6 increased, thus COX would increase, additional PGE2 could be incorporated into the GR. Ideally a measurement of PGE2 concentration made at the adrenal cortex would indicate this result.

Because of molecular similarity, competitive inhibition by PGE2 within the LBD will thereby influence the rate of cellular processing of CORT, which can be the source of dysfunctional cellular response, and hence systemic dysfunctional responses since GR is widely distributed, and that cortisol is linked to the hypothalamus, a site of autonomic control. Moreover as cortisol is also a permissive hormone, which itself impacts the processing rates of other hormones, competitive inhibition of cortisol would also alter the functional response of other homeostatic control processes. While this research has focused on the ligand binding domain of the glucocorticoid receptor, other receptors where cortisol acts could also be influenced by prostaglandins. This would include membrane bound receptors upon which cortisol may act, which would influence the potentiation of the cell. It has been noted that in addition to gene transcription through the LBD of GR, synaptic potentiation via cortisol has been examined [15, 16], which implies a non-genomic purpose as well, and thus these results pertain to non-genomic influences as well.

Based upon the correlations of disease and stress, the results of this study would seem more than coincidental that a conformer of PGE2, a biomarker of inflammation, and CORT, a biomarker of stress, would exhibit molecular similarity. With higher concentrations of PGE2 that are prevalent during states of disease, the signs and symptoms of disease would thus be amplified as the normative processing of cortisol would be altered due to competition for the GR. Moreover, therefore it is consistent that adrenal crisis due to cortisol insufficiency would result in flu-like symptoms, as the low rates of cortisol processing would be similar to that exhibited by competitive inhibition by PGE2. PGE2 is most notably arisen due to an infection, such as from gram-negative bacteria which possess lipopolysaccharides, and thus it would appear that many signs and symptoms of disease are due to the misprocessing of cortisol. Thus, the chronic nature of stress increases the likelihood that a form of illness will manifest; the dysfunctional processing of cortisol will amplify the signs and symptoms associated with prostaglandin concentrations due to disease. We believe these result to be of significant medical importance. Other experiments described in the research literature for which the developments of this paper can be evaluated include the study of [17] notes that chronic fatigue syndrome is correlated with low cortisol levels and HPA axis dysfunction, which is also cited in [18], noting that symptoms are enhanced with higher levels of stress. As indicated by this theory, hypocortisolism would be the result if PGE2 molecules occupying many LBD sites of the GR where cortisol nominally would reside, since the response of the hypothalamus will produce diminished levels of corticotropin releasing hormone because only a fraction of the GR responsible for its secretion are active, whereas the rest are ineffective due to interference of the competing PGE2 molecule. In addition, at the target site, the effects are diminished in actionable cortisol-GR associations thereby producing sub-normative trajectories. The inhibition of cortisol by prostaglandin at the LBD of GR would seem significant in the explanation and characterization of the primary source for dysfunction of the responses in signs and symptoms due to infectious agents. Research within this laboratory has linked the inhibition to responses inducing chills, fever, and other symptoms of infectious agents, including malaise or fatigue [10]. It is likely the source for a number of other symptoms of infectious disease and illnesses because of the wide distribution of the hormone cortisol and the prostaglandins.

## Supporting information

Supplemental to experimental analysis

