## Supplemental to experimental analysis for "Competitive Inhibition of Cortisol by Prostaglandins at the Ligand Binding Domain of Glucocorticoid Receptors"

February 4, 2020

### 1 Experimental Analysis

In [1], an experiment was conducted in which lipopolysaccharide (LPS) was injected and the response of the adrenal gland was measured over a 240 minute period, which included adrenocorticotropic hormone (ACTH), cortisol, TNF-$\alpha$, IL-1$\beta$, IL-6, and the phosphorylation of GR ($pGR$) to indicate its activity. In Spiga et al.'s analysis [1], a mathematical model was developed to characterize the assess the response of GR, as follows:

$$p\dot{G}R = k_{GR} f_{pGR}(ACORT) - \gamma_{pGR} pGR \tag{1}$$

in which the kinetics of phosphorylation are described as:

$$f_{pGR}(ACORT) = \frac{ACORT}{K_{ACORT} + ACORT} \tag{2}$$

Spiga and co-workers [1] fit the data to a model as presented in Figure 1

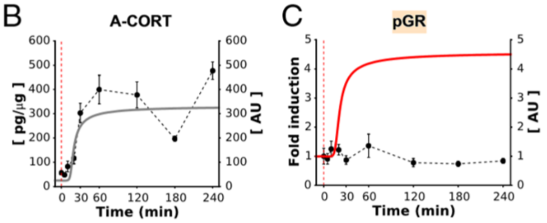


*Figure 1: Result from [1] indicating a lack of fitting of the model that assumed that cortisol would enable activity at the adrenal gland of GR due to a stimulus of ACTH via LPS.*

Because of the poor fit of $pGR$, the authors [1] indicated that the cytokine TNF-$\alpha$ was correlated to the observed response, and thus the model was updated as follows:

$$p\dot{G}R = k_{GR} f_{pGR}(ACORT)\phi_{pGR}(TNF\alpha) - \gamma_{pGR} pGR \tag{3}$$

where the additional factor was included in the modelling equations:

$$\phi_{pGR}(TNF\alpha) = \frac{K_{pGR}^{TNF\alpha}}{K_{pGR}^{TNF\alpha} + TNF\alpha} \tag{4}$$

The $TNF\alpha$ concentration was measured as shown in Figure 2

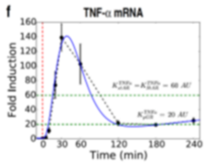


*Figure 2: A measurement of TNF-α taken when LPS was injected and the cytokines about the adrenal gland were measured from reference [1].*

which resulted in an improved fit as shown in Figure 3.

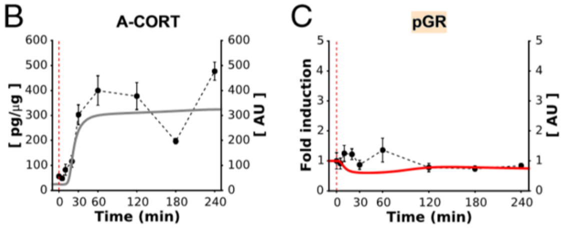


*Figure 3: An improved fit in [1] using the function $\phi_{pGR}(TNF\alpha)$ to quantitatively account from the lack of phosphorylation at GR after LPS injection even though cortisol concentrations increased.*

In this study, which utilizes the competitive inhibition of PGE2 on Cortisol at the GR, the model is instead configured as follows:

$$p\dot{G}R = k_{GR} f_{pGR}(ACORT, PGE2) - \gamma_{pGR} pGR \tag{5}$$

where the reaction rate kinetics are found from:

$$f_{pGR}(ACORT, PGE2) = \frac{ACORT}{K_{ACORT} + ACORT + K_I PGE2} \tag{6}$$

To determine the equivalent response of the data and of the modelled data, we have the following equivalency:

$$K_I PGE2 = K_{ACORT} + ACORT + \frac{K_{pGR}^{TNF\alpha}}{K_{pGR}^{TNF\alpha}} [K_{ACORT} + ACORT] \quad (7)$$

To obatin a numerical analysis the constants are taken as the same as [1] as the constants are $K_{pGR}^{TNF\alpha} = 20$ AU and $K_{ACORT} = 4.5$ AU, and the data is obtained from graphical results, and are listed in Table 1.

| Time (minutes) | experiment $ACORT$ (g/$\mu$g) | modeled $ACORT$ (pg/$\mu$g) | experiment $TNF\alpha$ fold induction | modeled $TNF\alpha$ fold induction |
|---|---|---|---|---|
| 0 | 20 | 0 | 0 | 0 |
| 5 | 30 | 0 | 0 | 0 |
| 10 | 80 | 0 | 12 | 12 |
| 20 | 120 | 100 | 70 | 70 |
| 30 | 300 | 260 | 140 | 140 |
| 60 | 400 | 280 | 100 | 100 |
| 120 | 350 | 300 | 20 | 20 |
| 180 | 200 | 305 | 20 | 20 |
| 240 | 480 | 310 | 20 | 20 |

*Table 1: Data from [1] used in the calcuation of this present study to determine the PGE2 concentration that lead to the competitive inhibition of the LBD at GR, as exhibited by a lack of GR activity at the adrenal gland.*

#### 1.1 Other Prostaglandins

In the family of prostaglandins, other configurations besides PGE2 show similarity in chemical affinity in comparison to CORT, and thus a capability for intermolecular activity within the LBD of GR and therefore competitive inhibition with CORT. Table 2 of the supplementary section lists several other prostaglandins, and indicates the main difference with respect to PGE2, and its possible difference in association with the LBD of GR under study. The result indicates that all prostaglandin have potential for competitive association with CORT, and thus are capable of altering the intended rate of gene expression of CORT.

| **Molecule** | **Formula** | **Difference from PGE2** | **Difference in Affinity at GR** |
|---|---|---|---|
| PGE1 | $C_{20}H_{34}O_5$ | C5-C6 single bond | $\Downarrow$ hydrophobic core |
| PGE3 | $C_{20}H_{30}O_5$ | double bond at C17-C18 | $\Downarrow$ C20 positioning |
| PGD2 | $C_{20}H_{32}O_5$ | OH at C9; O at C11 | $\Uparrow$ $I_{PGE2}$ hydrogen bonding<br>$\Downarrow$ $II_{PGE2}$ hydrogen bonding |
| PGF$\alpha$ | $C_{20}H_{34}O_5$ | OH at C9 | $\Uparrow$ $II_{PGE2}$ hydrogen bonding |

*Table 2: Analysis of the chemical affinity of other prostaglandins relative to PGE2 at GR, which to a first approximation would indicate that PGE1 and PGE3 would be slightly less stable within GR than PGE2, but both would should effective competitive inhibition. PGD2 shows improvements in PGE2 in hydrogen bonding as does PGF$\alpha$, which may result in slightly different kinetic effects than PGE2.*
